# Cholinergic modulation of membrane properties of calyx terminals in the vestibular periphery

**DOI:** 10.1101/2020.04.15.041491

**Authors:** Yugandhar Ramakrishna, Marco Manca, Soroush G. Sadeghi

**Author notes:** **Address of Correspondence:** Soroush G. Sadeghi, Center for Hearing and Deafness, 137 Cary Hall, 3435 Main St., Buffalo, NY 14214.

## Abstract

Vestibular nerve afferents are divided into regular and irregular groups based on the variability of interspike intervals in their resting discharge. Most afferents receive inputs from bouton terminals that contact type II hair cells as well as from calyx terminals that cover the basolateral walls of type I hair cells. Calyces have an abundance of different subtypes of KCNQ (Kv7) potassium channels and muscarinic acetylcholine receptors (mAChRs) and receive cholinergic efferent inputs from neurons in the brainstem. We investigated whether mAChRs affected membrane properties and firing patterns of calyx terminals through modulation of KCNQ channel activity. Patch clamp recordings were performed from calyx terminals in central regions of the cristae of the horizontal and anterior canals in 13 – 18 day old Sprague-Dawley rats. KCNQ mediated currents were observed as voltage sensitive currents with slow kinetics (activation and deactivation), resulting in spike frequency adaptation so that calyces at best fired a single action potential at the beginning of a depolarizing step. Activation of mAChRs by application of oxotremorine methiodide or inhibition of KCNQ channels by linopirdine dihydrochloride decreased voltage activated currents by ∼30%, decreased first spike latencies by ∼40%, decreased spike thresholds by ∼50%, and resulted in continuous firing during depolarizing steps. Interestingly, some of the calyces showed spontaneous discharge in the presence of these drugs. Together, these findings suggest that cholinergic efferents can modulate the response properties and encoding of head movements by afferents.

## INTRODUCTION

Vestibular sensors in the inner ear detect head movements and provide information that is carried by vestibular nerve afferents to the vestibular nuclei (VN) in the brainstem. The afferents are divided into regular (tonic) and irregular (phasic) groups based on the variability in interspike intervals of their resting discharges [1]. The most irregular afferents innervate the central regions of the sensory epithelium and have a specialized afferent terminal – the calyx – that covers the basolateral walls of one group of hair cells (type I) in adult amniotes [1, 2]. Interestingly, in squirrel monkeys and chinchillas, it has been shown that calyx terminals occasionally receive inputs from type II hair cells on their outer surface (with a 40:1 ratio of inner to outer surface ribbon inputs) [3]. The most regular afferents innervate the peripheral region of the epithelium by bouton terminals that contact another group of hair cells (type II). However, the majority of afferents innervate both type I and type II hair cells throughout the epithelium. The firing regularity of these ‘dimorphic’ afferents is correlated with the number of bouton terminals: those with fewer bouton terminals have higher irregular resting discharges [4, 5]. Irregular afferents innervating the semicircular canals have phasic properties and encode faster head movements, while their regular counterparts have tonic properties and provide detailed information about head movements, with lower detection thresholds [6]. Irregular afferents, as part of the phasic vestibular pathway (which also includes VN neurons) have been associated with vestibular compensation and adaptation [7–9].

Vestibular periphery of almost all vertebrates receives efferent inputs from neurons in the brainstem [10, 11]. In mammals, the net effect of efferent stimulation is an increase in resting discharge and a decrease in sensitivity of afferents, with a stronger effect on irregular afferents [11–13]. Both nicotinic (nAChR) and muscarinic (mAChR) acetylcholine receptors are expressed in the vestibular periphery [14–16] and mediate the fast [17] and slow components [18] of efferent mediated responses, respectively. Recent studies in knockout mice for CGRP [19] or α9 nAChR [20, 21] have shown defective VOR responses during high frequency rotations, suggesting that efferents are required for normal function of irregular afferents / phasic pathway. Furthermore, *in vitro* activation of mAChR in vestibular (Scarpa’s) ganglion neurons increases their firing rates during depolarization [22]. Systemic application of mAChR agonist *in vivo* also results in an increase in the response of vestibular nerve afferents [23].

In other brain areas, it has been shown that typically, the effect of mAChRs is mediated through inhibition of KCNQ potassium channels (i.e., M-currents) through a G-protein mediated pathway [24–26]. KCNQ potassium channels are one of the major contributors to the conductances observed in calyx terminals and play a role in setting the regularity of the resting discharge in vestibular ganglion neurons [27–35]. Four of the five KCNQ subtypes are expressed in the nervous system (KCNQ2 – KCNQ5 or Kv7.2 – Kv7.5) [24–26, 36] and all four are also present in the vestibular periphery [31, 37, 38]. Modulation of KCNQ channels have been shown to affect neuronal response properties in different brain areas: vestibular ganglion neuron somas [22, 27], sympathetic neurons [39], dorsal root ganglia [40, 41], sensory Aδ and C-fiber peripheral endings [42, 43], peripheral baroreceptor afferents [44], and CA1 pyramidal neurons [45, 46]. Decrease in the current and / or activation and deactivation rates of KCNQ channels result in a decrease in spike frequency adaptation and an increase in excitability of neurons, a decrease in their firing threshold, and induction of spontaneous discharge (mainly attributed to the expression of KCNQ channels near the axonal spike generation zone) [47–51].

While the above previous studies on the function of mAChR and KCNQ channels in the vestibular periphery are informative, they also have some limitations or may differ from those in mammals. First, these studies have been performed on the soma of ganglion cells, which do not receive cholinergic efferent inputs and the effect of drugs in this area is an indirect indication of efferent-mediated effects on afferent terminals. Second, most ganglion cell receives inputs from many calyx and bouton terminals, which might be differentially affected by mAChRs activation. Finally, systemic application of KCNQ antagonists also affects K^+^ recycling in the inner ear [36], which can indirectly affect the vestibular function. One study that shows a link between mAChR and slow efferent-mediated changes in afferent firing rate has been performed in turtle [18]. Recordings from a group of turtle vestibular afferents that only have calyx terminals was used for this purpose. However, as mentioned above, most afferents in mammals receive inputs from both type I and type II hair cells and their properties therefore might differ from those in the turtle. In the present study, we used patch clamp recordings from calyx terminals to directly investigate whether mAChR activation affected their membrane properties in rats. We found that application of oxotremorine methiodide (oxo-M), a mAChR agonist resulted in attenuation of voltage sensitive currents and enhancement of excitatory membrane properties, similar to what has been observed by inhibition of KCNQ channels. Additionally, the oxo-M effect could be completely blocked by KCNQ channel antagonists. Typically, under *in vitro* conditions, most calyces in the central regions of the neuroepithelium display no spontaneous firing and only fire a single action potential during step depolarizations [52, 53]. In the presence of mAChR agonist, calyces fired many action potentials during step depolarizations, most likely due to inhibition of KCNQ channels and decreased spike frequency adaptation. Interestingly, some of the calyces even developed spontaneous activity in the presence of mAChR agonist. In addition, calyces fired action potentials in response to smaller step depolarizations (i.e., decreased AP threshold), with decreased first spike latencies. Similar results were observed with application of KCNQ antagonist linopirdine dihydrochloride. Our findings suggest that without mAChR activity, calyces cannot respond to stimuli due to open KCNQ channels and the resultant high spike frequency adaptation, a finding supported by our previous *in vivo* findings [54]. We propose that through activation of mAChRs, cholinergic efferent inputs provide the means for changing membrane properties of calyx terminals and consequently, spontaneous activity and response properties of irregular afferents.

## MATERIALS AND METHODS

### Animals

All procedures were approved by the Institutional Animal Care and Use Committee of the State University of New York at Buffalo, Buffalo, NY and the Johns Hopkins University, Baltimore, MD, and carried out in strict accordance with the recommendations in the Guide for the Care and Use of Laboratory Animals of the National Institute of Health. Fifty seven Sprague-Dawley rats (Charles River Laboratories), 13 to 18 day old and of either sex were used for the experiments.

### Tissue preparation

Dissection of the end organ and tissue preparation were performed as described previously [52]. Briefly, rats were deeply anesthetized by isoflurane inhalation and decapitated. The bony labyrinth was then removed and placed in extracellular solution. Under the microscope, the bone was opened over the ampullae of the horizontal and anterior canals and the utricle. The membranous labyrinth containing the horizontal canal, anterior canal, the utricle, and the Scarpa’s ganglion was taken out of the bone. The top of the membranous labyrinth was opened to expose the neuroepithelium of the two cristae and the utricle. It is important to note that while efferent terminals are present in this preparation, efferent fibers are cut and separated from their somata located in the brainstem[52, 53]. These efferent fibers and terminals can be stimulated by electrical or optogenetic stimulation [55]. Note that while the drugs applied in the current study can potentially affect these efferent terminals, this will not affect our conclusions. Furthermore, there is currently no evidence that mAChR exist on efferent terminals.

### Electrophysiology recordings

The preparation was secured under a pin on a coverslip, transferred to the recording chamber, and perfused with extracellular solution at 1.5 −3 ml/min. The extracellular solution contained (in mM): 5.8 KCl, 144 NaCl, 0.9 MgCl_2_, 1.3 CaCl_2_, 0.7 NaH_2_PO4, 5.6 glucose, 10 HEPES, 300 mOsm, pH 7.4 (NaOH). The intracellular solution contained (in mM): 20 KCl, 110 K-methanesulfonate, 0.1 CaCl_2_, 5 EGTA, 5 HEPES, 5 Na_2_ phosphocreatine, 4 MgATP, 0.3 Tris-GTP, 290 mOsm, pH 7.2 (KOH). To perform patch-clamp recordings, tissue was visualized with a 40x water-immersion objective, differential interference contrast (DIC) optics (Examiner D1 Zeiss microscope), and viewed on a monitor via a video camera (optiMOS, Qimaging). Patch-clamp recording pipettes were fabricated from 1 mm inner diameter borosilicate glass capillaries (World Precision Instruments, item# 1B100F-4). Pipettes were pulled with a multistep horizontal puller (P-1000, Sutter), coated with Sylgard 184 (Dow Corning) and fire polished. Pipettes were filled with the internal solution and at the beginning of each experiment, resistances were measured in voltage clamp during a depolarizing 10 mV step (from −79 mV) with the tip of the electrode submerged in the extracellular solution. Electrodes typically had resistances of 5-10 MΩ. All recordings were performed at 23 – 25 °C (room temperature). Application of drug solutions was performed (VC-6 channel valve controller, Warner Instruments, Hamden, CT) using a gravity-driven flow pipette (∼100 μm tip diameter) placed near the recorded calyx (∼200 μm). The above solutions result in a liquid junction potential of −9 mV [52], which is corrected for voltages presented in figures and text. Drugs were dissolved daily in the extracellular solution to their final concentrations from frozen stocks. Agonist of mAchR, Oxotremorine-M (Oxo-M) (20 μM) and KCNQ antagonist Linopirdine (20 μM) [56] were purchased from Tocris. We used concentrations higher than the usual 10 μM for Linopirdine [27, 37] and Oxo-M [22] in order to get larger effects more consistently in larger number of recordings.

### Experiment protocol

Whole cell patch clamp recordings were performed from calyx afferent terminals in the central region of the cristae of either anterior or horizontal canals. All measurements were acquired using pCLAMP 10 software in conjunction with a Multiclamp 700B amplifier (Molecular Devices), digitized at 50 kHz with a Digidata 1440A, and filtered at 10 kHz.

Calyces were identified as thickenings around type I hair cells, which were identified by their typical morphology [52]. Membrane capacitance and series resistance were electronically compensated during recordings (correction and prediction circuits set to 75 – 80% with a bandwidth of 10 – 15 kHz). To have a uniform initial condition, cells were held at a holding potential of −79 mV, which was slightly hyperpolarized compared to rest and resulted in holding currents of < 200 pA (on average: 127 ± 15 pA). Since our experiments lasted ∼20 min and whole cell currents can change over time [37], all data were collected 5-7 min after the membrane was ruptured for whole cell recording to minimize time dependent changes. For drug applications, data were collected 9-12 min after start of application to have consistently stable, significant and reliable effects (see Results and Fig. 4).

During voltage clamp recordings, a voltage step protocol was used (Fig. 1A), which consisted of an initial holding potential of −79 mV (100 ms), a hyperpolarizing step to −129 mV (100 ms), 20 mV steps between −129 and +11 mV (300 ms), and return to holding potential of −79 mV. As previously described [37, 52, 53, 58], calyx recordings exhibited (1) large Na^+^ inward currents corresponding to action potentials, at the beginning of depolarizing steps (starting at −69 mV or −49 mV; average: −62.6 ± 3.3 mV) with amplitudes of several nA, (2) delayed rectifier type outward currents during depolarizing voltage steps (mainly due to KCNQ activation), (3) slowly deactivating currents during the initial hyperpolarizing step (mainly due to KCNQ deactivation), and (4) hyperpolarization-activated (I_h_) currents during hyperpolarizing steps.

**Figure 1.**
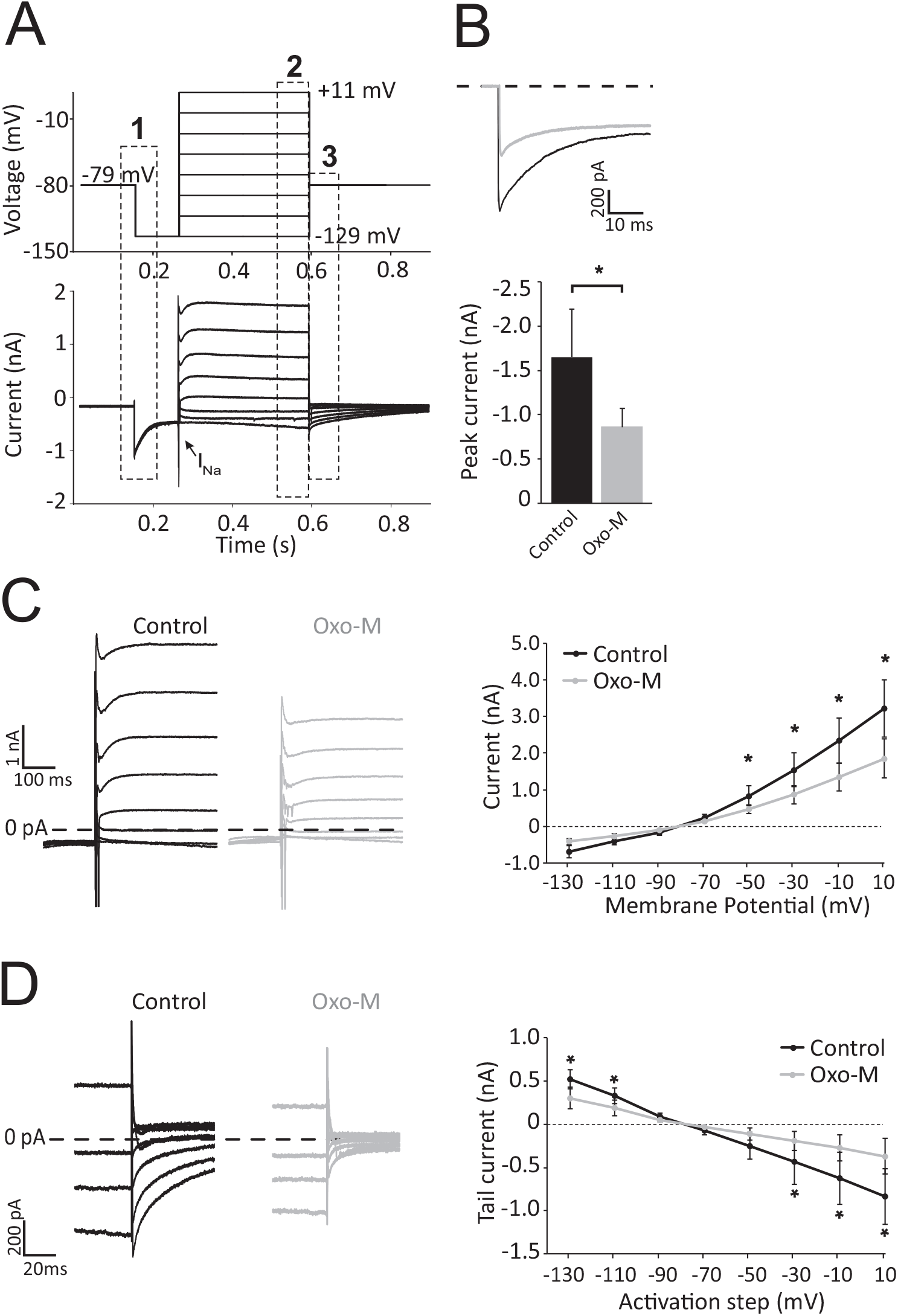
Activation of mAChR attenuates voltage activated currents. (A) Response of a calyx to the voltage step protocol (holding potential of −79 mV, initial hyperpolarizing step to −129 mV, 20 mV steps between −129 mV and +11 mV, and return to −79 mV after the steps). Calyx recordings exhibited slowly deactivating currents during initial hyperpolarizing steps (dashed box 1), delayed rectifier type outward currents during depolarizing voltage steps (dashed box 2), and tail currents (dashed box 3). I_Na_: sodium inward currents corresponding to action potential at the beginning of depolarizing steps. (B) Deactivating currents during the initial hyperpolarization decreased in the presence of oxo-M (20 μM). Top traces are examples of responses in control condition (black) and in the presence of oxo-M (grey). Bottom bar graphs show a significant decrease in average responses (n = 9 calyces). (C) Outward currents produced by step protocol decreased during oxo-M application in an example recording (left). Average responses (right) showed a significant decrease for depolarizing steps above resting membrane potential. (D) The amplitude of tail currents after the steps decreased during oxo-M application in an example recording (left). Average tail currents (right) significantly decreased after steps that were > 20 mV away from the resting membrane potential. In all panels significant changes are marked by asterisks. Note, all changes in average currents are presented relative to the current at the initial holding potential of −79 mV.

During current clamp recordings, the membrane potential was first measured with no current injection (i.e., resting membrane potential). Current injection was adjusted to hold the membrane potential between −69 mV to −79 mV for 10 s. None of the recorded calyces in the present study had spontaneous firing at the resting potential or at ∼ −79 mV. Calyces were then depolarized by injecting step currents of 100 pA, 200 pA, 300 pA, 400 pA, and 500 pA. The steps were typically about 2 s in duration, however, we sometimes waited up to 5 s in order to get closer to a plateau potential. A wait time of ∼5 s was required between different steps so that the membrane potential returned to the initial potential.

### Data analysis

Clampfit 10 (Molecular Devices), minianalysis (Synaptosoft), and Graphpad Prism (version 5) software was used for the analysis of data. Results are reported as mean ± SE. Paired t-test was used for comparison between two parameters and repeated measures two way ANOVA with Bonferroni post hoc test was used for comparisons of more than two conditions. Level of statistical significance was set at α = 0.05.

To study the effect of various drugs in voltage clamp, currents were measured at 3 different parts of the voltage step protocol (Fig. 1A, dashed boxes 1 – 3): (1) the peak amplitude of the inactivating current during the initial hyperpolarization step from −79 mV to −129 mV, (2) current amplitude over the last 50 ms of the steps (from −129 mV to +11 mV), and (3) peak amplitude of tail currents at −79 mV during the first 50 ms after steps. For tail currents, peak amplitudes were plotted against holding potential at the preceding step (‘activation step’ in figures). All changes in currents were calculated relative to the current at the initial holding potential of −79 mV.

During current clamp steps, the smallest of the five steps that generated an AP was considered as the threshold. If a spike was generated with 100 pA step, smaller current steps were also injected to find a value closer to the actual threshold. The latency of the first spike was measured as the time from the beginning of the step to the peak of first AP for this threshold step. Finally, the number of APs during the 2 s steps were also counted. Results were compared before and after application of drugs.

## RESULTS

Patch clamp recordings were performed as described before [52] from the calyces in the central zone of the cristae of the horizontal or anterior canals in Sprague-Dawley rats. Thirteen to 18 day old animals were used (n = 57) so that type I hair cells and calyx afferent terminals had acquired their characteristic morphological and electrophysiological properties [37, 59–61]. The average resting membrane potential of calyces during whole-cell recordings was −71.9 ± 3.8 mV (n = 27), comparable to numbers reported previously [52, 53, 62]. A voltage step protocol was used to study the voltage sensitive KCNQ potassium currents (Fig. 1A). During the initial hyperpolarization, calyx recordings showed an early slowly deactivating current, mainly attributed to KCNQ channels [37, 52, 53, 59] followed by a hyperpolarization-activated (I_h_) current [29, 52, 58]. During depolarizing steps, an inward Na^+^ current (corresponding to an action potential) was observed at the beginning of steps and later slow outward currents most likely due to activation of different K^+^ channels (Sadeghi et al. 2014) including (1) delayed rectifier type currents by KCNQ channels [37], (2) voltage sensitive BK channels [30], and (3) calcium sensitive SK channels [63]. No spontaneous (resting) discharge was present in any of our control recordings from the central regions of the cristae.

### Voltage sensitive currents were suppressed by mAChR activation

To investigate whether mAChR receptors were present on calyx terminals and their possible effect on membrane properties of calyces, we applied the mAChR agonist oxo-M (20 μM) during patch clamp recording from calyces. For all recordings during oxo-M application (n = 9), a decrease in voltage sensitive currents was observed, suggesting the presence of mAChR receptors. In the presence of oxo-M, the amplitude of the inward current during the initial hyperpolarizing step (Fig. 1B) decreased from −1649.89 ± 538.67 pA to −858.9 ± 215.6 pA (paired t-test, p = 0.0006). It has been shown that this inward current is mainly due to slow deactivation of KCNQ channels [37], suggesting that mAChR activation by oxo-M inhibited the KCNQ current. Oxo-M application also decreased voltage activated currents during depolarizing steps (Fig. 1C). For the population of recorded calyx terminals, application of oxo-M resulted in an inhibition of currents in response to depolarizations above resting membrane potential (−49 mV to +11 mV steps) (repeated measures two way ANOVA: p < 0.0001, Bonferroni posthoc test: p > 0.05 for steps ≤ −69 mV, p < 0.05 at −49 mV and p < 0.001 for steps > −49 mV). Finally, tail currents recorded when membrane potential returned to −79 mV at the end of steps were also decreased during oxo-M application (Fig. 1D). Tail currents are most likely due to increased extracellular [K^+^] due to outward voltage sensitive K+ currents during depolarizing steps.. Based on the observed decrease in voltage activated K^+^ currents during depolarizing steps with oxo-M application, decreases in tail currents were also expected. While the small tail currents were not affected significantly, the amplitude decreased after all other steps (Fig. 1D, asterisks) (repeated measures two way ANOVA, p = 0.0001, Bonferroni posthoc, p < 0.05 for steps < −109 mV, p > 0.05 for steps −89 mV to −49 mV and p < 0.001 for steps > −49 mV for comparisons at each voltage before and after oxo-M). Note, voltages that produced small tail currents close to K+ reversal potential in the control condition will be minimally affected by linopirdine. Together, these results confirm that mAChRs are expressed by calyces and that their activation inhibits voltage sensitive K^+^ currents.

### Suppression of voltage sensitive currents by KCNQ channel inhibition

We next investigated the effect of inhibition of KCNQ channels on calyx membrane properties. We first applied the KCNQ antagonist linopirdine (20 μM) during the same step protocol as described above. In the presence of linopirdine, inward and outward current changes were almost identical to that observed with oxo-M application. The inward current during the initial hyperpolarizing step was reduced in the presence of linopirdine (Fig. 2A) and on average (n = 6) decreased from −966 ± 255 pA in co580 ± 89 pA after linopirdine application (paired t-test, p = 0.017). For depolarizing steps, linopirdine reduced the outward K^+^ currents (Fig. 2B) during depolarizing steps of −49 mV to +11 mV (repeated measures two way ANOVA, p < 0.0001, Bonferroni posthoc, p > 0.05 for steps ≤ −69 mV, p < 0.05 for −49 mV step and p < 0.001 for steps > −49 mV). Finally, while linopirdine had no effect on tail currents after steps of −69 mV and −89 mV (i.e., near K^+^ reversal potential), it reduced the outward and inward tail currents after other steps (Fig. 2C, asterisks) (repeated measures two way ANOVA, p < 0.0001, Bonferroni posthoc, p < 0.01). The similarities between these results and those obtained with oxo-M application, strongly support the idea that oxo-M effects were mediated by the interaction between mAChR and KCNQ channels, similar to that shown in the vestibular ganglion [22] and neurons in other brain areas [25]. To directly show that the mAChR effect in the calyx is mediated through KCNQ channels, in a subset of recordings, linopirdine was applied initially to suppress KCNQ-mediated currents, followed by oxo-M (plus linopirdine) application (Fig. 3). As expected, linopirdine decreased voltage sensitive K^+^ currents during depolarizing steps (Fig. 3A) of −129 mV to +11 mV. Application of Oxo-M (with linopirdine) did not have any further effect on K^+^ currents. For 5 calyces recorded, there were significant decreases in K^+^ currents at the two highest depolarizations after linopirdine application (Fig. 3B, repeated measures two way ANOVA, Bonferroni post-hoc test, p < 0.01), with no further decrease when oxo-M was added. This result shows that the oxo-M effect is mediated through KCNQ channels in the calyx afferent terminal.

**Figure 2.**
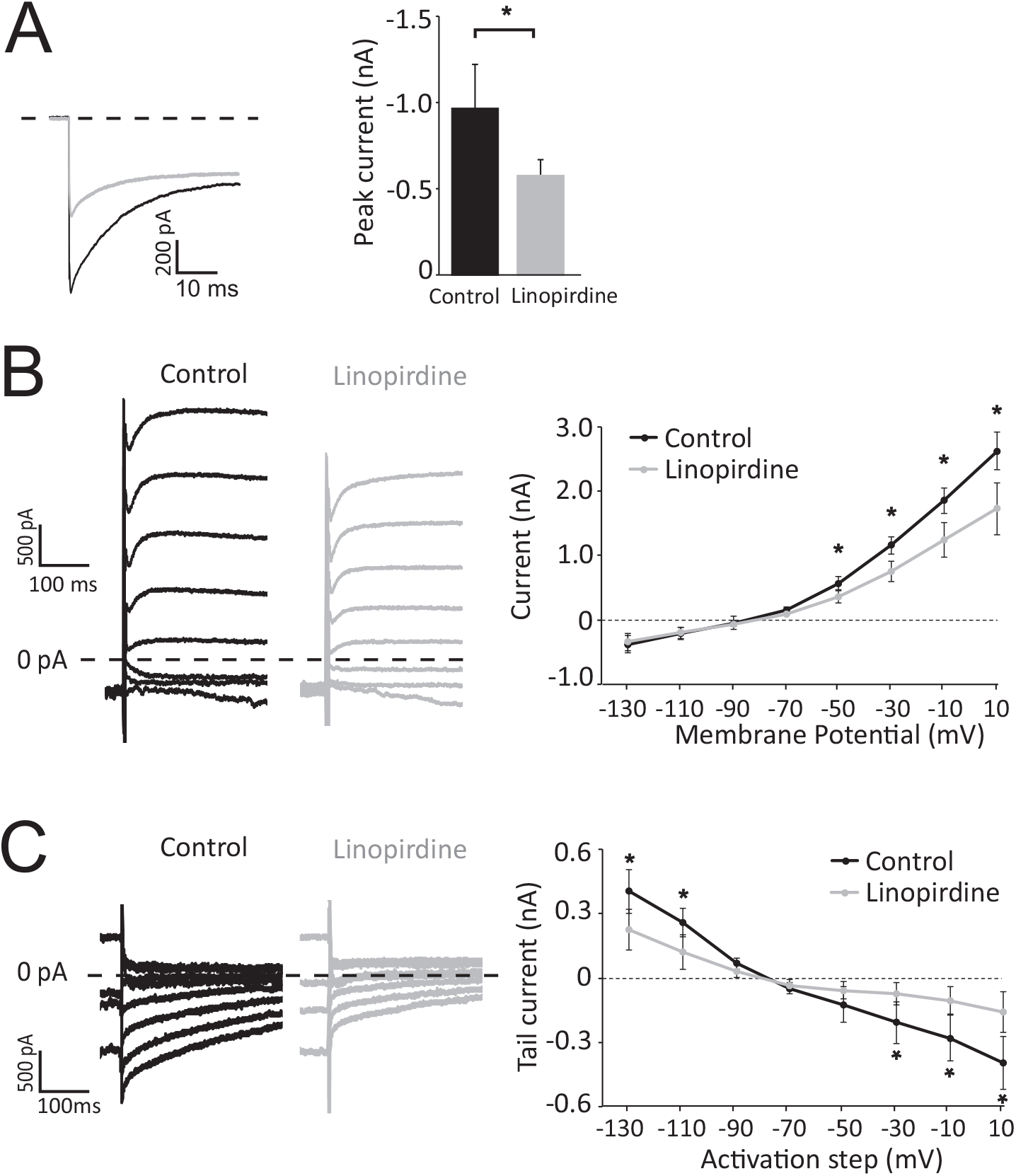
Inhibition of KCNQ channels attenuates voltage activated currents. (A) Deactivating currents during the initial hyperpolarization decreased in the presence of linopirdine (20 μM). Traces on the left show examples of responses under control condition (black) and during linopirdine application (grey). Linopirdine resulted in a significant decrease in average responses (n = 6 calyces). (B) Outward currents produced by step protocol decreased during linopirdine application in an example recording (left). Average responses (right) showed a significant decrease for depolarizing steps above resting membrane potential. (C) Tail currents produced after steps decreased during linopirdine application in an example recording (left). Average tail currents (right) were significantly decreased after steps that were > 11 mV away from the resting membrane potential. In all panels significant changes are marked by asterisks. Note, all changes in average currents are presented relative to the current at the initial holding potential of −79 mV.

**Figure 3.**
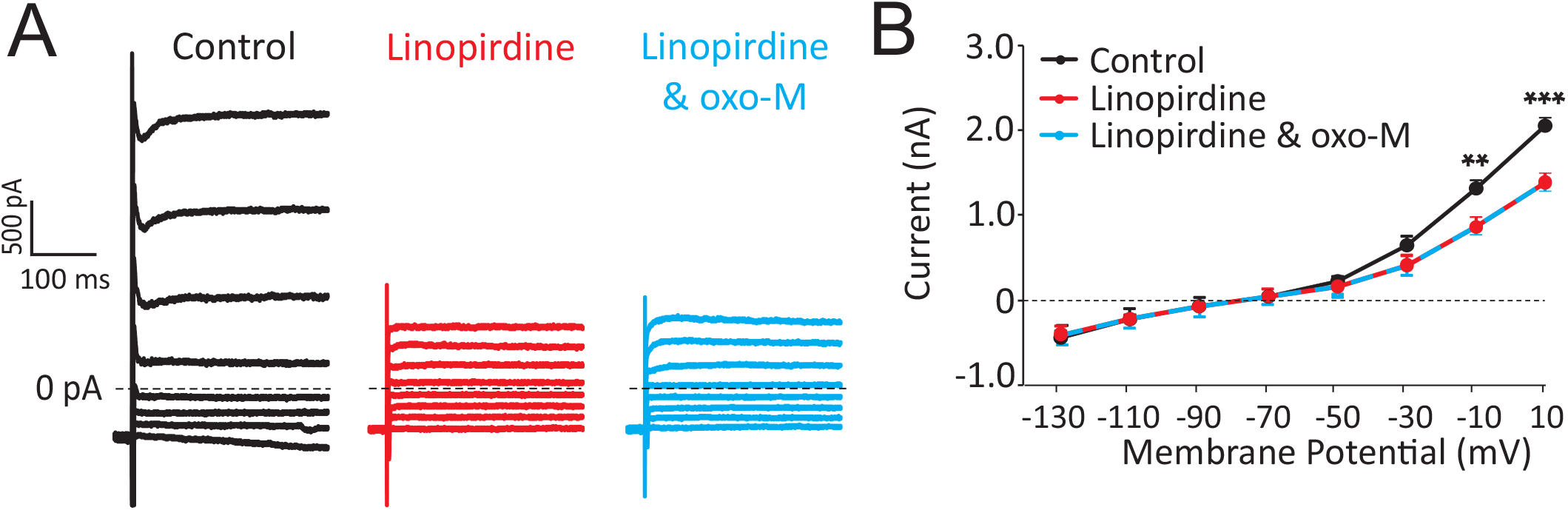
Effect of mAChR is mediated through KCNQ channels. (A) Example calyx recording showing that application of the KCNQ antagonist linopirdine decreased voltage sensitive currents and blocked the effect of mAChR activation by oxo-M. (B) Average responses (right) showed a significant decrease in currents in the presence of linopirdine and no further change when oxo-M was added, suggesting that mAChR effect was mediated through KCNQ channels. ** and *** represent p<0.01 and p<0.001, respectively.

### The effect of duration of recordings and drug application

We noticed early during experiments that both linopirdine and oxo-M typically required about 3 – 5 min of application before any effect was observed and the effects increased with time. Similar long wait times had previously been reported for other metabotropic agonists applied to the cochlea [67] and for KCNQ antagonists used in the hippocampus [46]. However, previous studies had suggested that currents obtained by intracellular patch clamp recording could be affected by duration of the recording [37]. Since this could have been a source of variability in our results, we investigated possible effects of long wait times on the recorded currents. In a set of experiments, we kept recordings for up to 30 min or more and compared responses at different time points. In the control condition, we did not observe any significant change in K^+^ currents up to 15 min (n = 25) or even 30 min (n = 7) after the beginning of intracellular recording (Fig. 4A). This result suggested that changes in patch clamp recording itself were not a major contributor to the observed effects of drug applications. During linopirdine application (n = 6), a decrease in K^+^ currents was observed after 3 – 5 min and reached significant levels by 7 – 9 min (Fig. 4B). The effect of oxo-M (n = 9) also became significant 7-9 min after starting the application (Fig. 4C) and further recordings up to 12 min did not show further change in its effect. To be uniform in our conditions and to decrease the variability of drug effects, all our data were collected 9-11 min after application of linopirdine or oxo-M.

**Figure 4.**
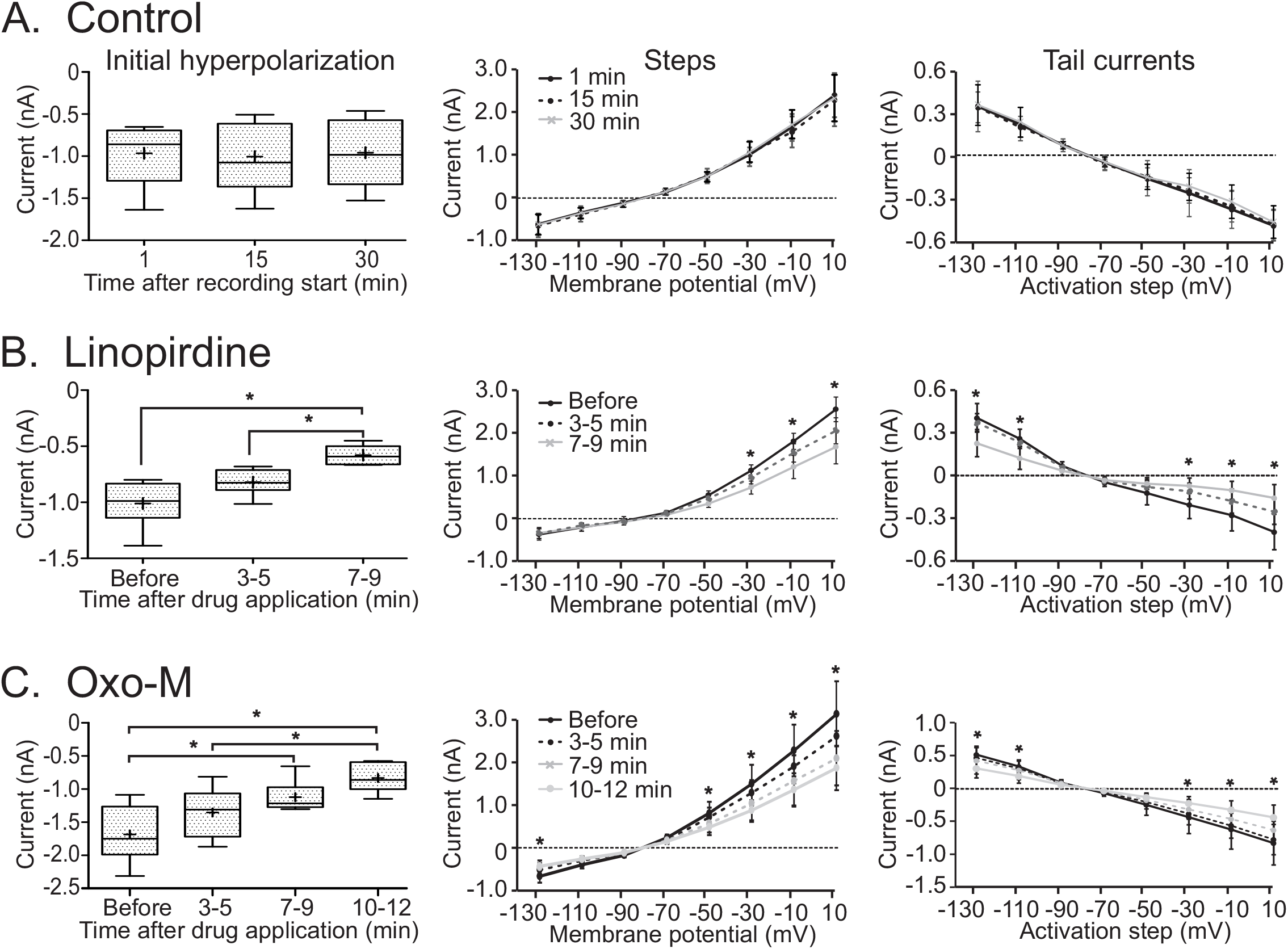
Effect of duration of patch clamp recording and drug applications on calyx responses. (A) For the control condition, the currents did not show any significant change over time, even after 30 min of recording. (B) The effect of linopirdine application increased over time, reaching significant values after 7-9 min of application. (C) The effect of oxo-M increased over time, reaching significant values of inhibition after 7-9 min of application. The effect did not increase further when measured after 10-12 min of drug application. In all panels significant changes are marked by asterisks; details are provided in the Results. Box plots show the mean (plus sign), median (line), 25 – 75% interval (box), and 10 – 90% interval (whiskers). Note, all changes in average currents are presented relative to the current at the initial holding potential of −79 mV.

### Activation of mAChRs increased the sensitivity of calyx terminals

We next used current clamp recordings to investigate changes in firing properties and more functional aspects of mAChR activation. To have a similar initial condition for all recordings, we injected currents to keep the initial membrane potential between −60 mV to −70 mV. We then injected current steps of 100 – 500 pA in 100 pA steps (Fig. 5A, top row) to depolarize the calyx. In control condition, none of the recorded calyces (n = 15) showed spontaneous firing and the threshold for generating an action potentials (AP) was reached by injecting an average current of 318 ± 135 pA. In control, some calyces did not fire any AP even with the largest depolarization (Figs. 5A1 and 5A2). For the rest of the recordings, calyces generated only a single AP at threshold as well as at higher membrane potentials, consistent with a previous report [53]. Some calyces displayed oscillations in membrane potential after firing an AP, similar to that shown in some of Scarpa’s ganglion neurons [27]. The latency of the single spike was 5 ± 0.36 ms at threshold and decreased as a function of increasing amplitudes of injected currents / depolarizations.

**Figure 5.**
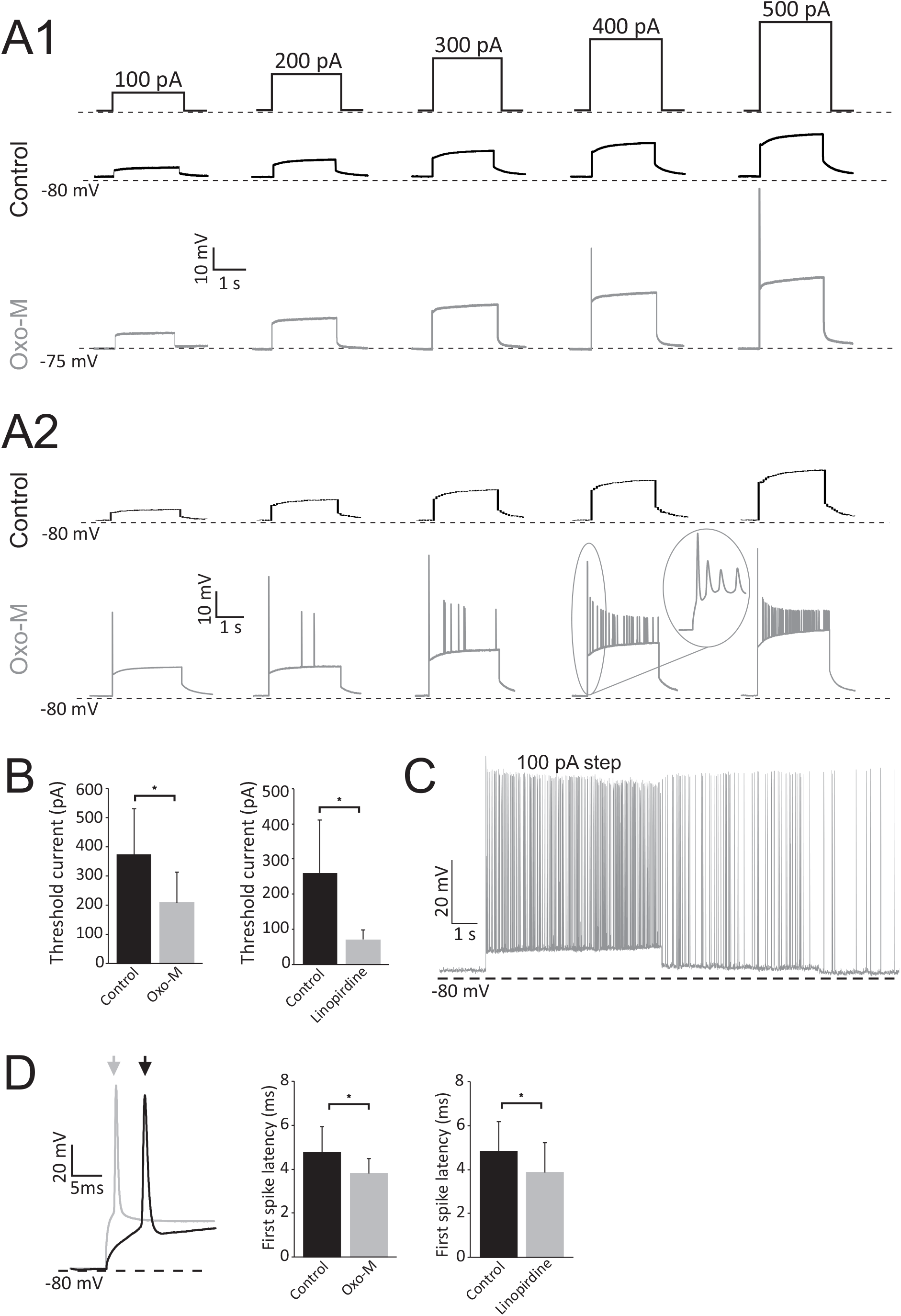
Calyces become more sensitive and respond faster following activation of mAChR or inhibition of KCNQ channels (oxo-M: n = 9, linopirdine: n = 6). (A) Examples of current clamp recordings from calyces that did not fire any APs in the control condition in response to injection of 100 – 500 pA current steps, but in the presence of oxo-M fired a single AP at the beginning of a step (A1) or many APs during the step (A2). Similar results were observed during linopirdine application (not shown). (B) The amplitude of the current required for AP generation (i.e., current threshold) decreased significantly during oxo-M or linopirdine application. Note that in the presence of oxo-M, currents below 100 pA were also tested to find a closer value to the actual threshold. (C) Some calyces showed spontaneous AP firing in the presence of oxo-M application. Similar results were also observed during linopirdine application (not shown). (D) The latency of the first spike decreased during application of oxo-M (grey arrow) or linopirdine (not shown) compared to control condition (black arrow). Although small, the decrease was significant. In all panels significant changes are marked by asterisks.

After oxo-M (n = 9) or linopirdine (n = 6) application, calyces became more excitable. As mentioned above, all values presented here were recorded 9 – 11 minutes after beginning of drug application. Most calyces (5/9 with oxo-M and 3/6 with linopirdine) generated a higher number of spikes after oxo-M or linopirdine application compared to control conditions (Figs. 5A1 and 5A2), particularly for 300 pA, 400 pA, and 500 pA steps. Furthermore, the minimum current required for AP generation (i.e., threshold current) significantly decreased (Fig. 5B) from 372 ± 156 pA to 211 ± 102 pA with oxo-M (paired t-test, p=0.03) and from 260 ± 151 pA to 70 ± 27 pA with linopirdine (paired test, p = 0.03). Finally, 3 of the 15 recorded calyces exhibited spontaneous firing in the presence of oxo-M or linopirdine, which occurred either after a current step (Fig. 5C) or spontaneously in between trials. Induction of spontaneous activity following KCNQ inhibition has been reported in other brain areas and is attributed to inhibition of KCNQ channels in the vicinity of the spike generation zone [25, 48–50]. Such colocalization is also observed in the calyx [31] and could contribute to the observed increase in spike generation with inhibition of KCNQ channels in the calyx.

During drug applications responses also became faster in that the first spike latency decreased during depolarization. Figure 5D shows the decrease in average threshold currents during application of oxo-M (4.8 ± 1.15 ms vs. 3.8 ± 0.67 ms, paired t-test, p=0.033) or linopirdine (4.8 ± 1.37 ms vs. 3.9 ± 1.35 ms, paired t-test, p=0.036). Similar to the control condition, the latency of the first spike decreased with increasing amplitude of step currents so that the first spike latency remained lower after drug applications even for 500 pA steps (paired t-test, p<0.05).

## DISCUSSION

The results of the present study suggest an important role for the cholinergic efferent vestibular pathway in shaping membrane properties and spike frequency adaptation of calyx terminals. We show that *in vitro*, calyx terminals in the central region of the cristae at best fire a single AP during step depolarizations. With activation of mAChRs, calyces fire more APs during such steps (i.e., increased sensitivity), which is also associated with a decrease in the minimum current required for AP generation (i.e., decreased current threshold) and a decrease in the latency of the first spike during steps (i.e., faster response). Interestingly, mAChR activation results in spontaneous firing in some calyces. These effects are due to a decrease in voltage sensitive K^+^ currents. Inhibiting KCNQ channels blocked the effect of mAChR agonists, suggesting that these receptors function through inhibition of KCNQ channels similar to other brain areas. Together, these results show that efferent inputs and activation of mAChRs can change membrane properties of calyces through inhibition of KCNQ channels. We propose that such cholinergic modulation of calyx membrane properties provides the means for continuous adjustment of the sensitivity and response dynamics of calyx terminals and by extension vestibular afferents, particularly irregular fibers.

### Modulation of subthreshold potentials by efferents: effects on quantal and non-quantal transmission

We have recently shown that inhibition of efferents results in a decrease in resting discharge of afferents and even complete shutdown of the most irregular afferents [54]. The present study supports these previous findings. The *in vitro* tissue preparation consists of the anterior and horizontal canal cristae and utricular macula along with afferent cell bodies in the Scarpa’s ganglion [52, 53], but it is devoid of efferent cell bodies that are located in the brainstem. Lack of spontaneous efferent inputs in this preparation was previously shown by block of all spontaneous synaptic events in calyx recordings in the presence of glutamate receptor antagonists [52]. We propose that the activity of different types of K^+^ channels in calyces [31, 37, 63, 70], particularly KCNQ channels that are open near the resting membrane potential results in spike frequency adaptation, a property shared with other brain areas [25]. In particular, KCNQ channels are unique in that they become inactive only at very hyperpolarized potentials and are still open close to the resting membrane potential. Therefore, they can play an important role in modulating subthreshold membrane properties. KCNQ inhibition can also make neurons more responsive to synaptic inputs (i.e., larger EPSPs with the same presynaptic condition) [71]. The type I hair cell – calyx synapse is unique in that in addition to the typical quantal signal transmission, it also provides a non-quantal component that is mediated by accumulation of K^+^, glutamate, and H^+^ in the closed synaptic space between the calyx and type I hair cell [52, 62, 64, 72–75]. The non-quantal transmission has been proposed as a mechanism for encoding fast movements [53]. Modification of the activity of KCNQ channels by mAChR provides the means for adjusting the firing properties of calyx terminals, which can affect their responses to both quantal (through both type I and type II hair cells) and non-quantal signal transmission.

### Efferent inputs can affect the response properties of dimorphic afferents

Of the two groups of afferents, irregular fibers have phasic properties and encode fast head movements, while regular fibers have tonic properties and carry detailed information about head movements by the semicircular canals and changes in the gravity vector (i.e., head tilt) by otoliths [6, 76]. Most afferents have ‘dimorphic’ terminals and receive inputs from both type I and type II hair cells [1]. Irregular dimorphic afferents have fewer bouton terminals than regular dimorphic afferents [4, 5], suggesting that the ratio of calyx-to-bouton terminals affects afferent properties. Note that some of the input to dimorphic afferents could come from type II hair cells by direct synapses on the outer surface of calyces [3, 77]. Our results provide evidence that one way for changing the contribution of phasic / tonic inputs in dimorphic afferents is through changes in KCNQ channel activity by cholinergic efferent inputs, which affects the relative contribution of inputs from calyces to these fibers.

Our results show that cholinergic efferent inputs on calyces in the central region could play an important role in modifying response properties of calyx membrane properties. However, it should be noted that there are differences in membrane properties of hair cells and afferent terminals between the central and peripheral regions of the neuroepithelium [31, 32, 53, 72]. In the present study, we only recorded from calyces in the central regions of the cristae and cannot speculate about the effects on calyces that innervate the peripheral regions. Afferents that innervate the central region have the most irregular resting discharge [1], receive inputs only from calyx terminals or have fewer bouton terminals compared to peripheral regular dimorphic afferents [4, 5], show the largest efferent-mediated effects [11–13, 54], and have an abundance of KCNQ channels [31].

### Effect of mAChR activation on spike generation by calyx terminals

With both oxo-M and linopirdine, the first spike latency was reduced compared to the control condition. The importance of first spike latency has been studied in transmission of sensory information and different stimulus features in several sensory systems, including the auditory [79, 80], visual [81], and somatosensory [82] systems. In the vestibular periphery, it is possible that the first spike carries information about very fast head movements when afferents mostly phase lock to the stimulus [83]. Otolith afferents are particularly prone to such phase locking [76, 84]; a response that could underlie short latency vestibular sensory evoked potential (VsEP) responses measured by subcutaneous electrodes [85]. A recent *in vivo* study has shown supporting evidence for an enhancement of VsEP responses after systemic application of KCNQ antagonist XE-991 [23]. Future *in vivo* studies are required to address whether first spikes play a role in transmitting information about specific aspects of head movement.

Both oxo-M and linopirdine strongly enhance firing of action potentials by calyx terminals, similar to that shown in dissociated ganglion neurons in response to application of oxo-M or KCNQ antagonists [22, 27, 33] and in support of *in vivo* responses after systemic application of KCNQ antagonist XE-991 [23]. These excitatory responses are in contrast to the hyperpolarizing effect of mAChR activation on type II HC [86–88]. The latter has been attributed to activation of the M2 subtype of mAChR and its effect on opening of calcium channels and subsequently, big current (BK) potassium channels [89]. While all subtypes of mAChR are expressed in the vestibular sensory epithelium [16], the excitatory responses of calyces suggest the presence of a different subtypes of mAChR, most likely M1, M3, or M5 that are known to modulate KCNQ channels [25, 90] in calyx afferent terminals. Such peripheral changes are most likely part of the effect of nonspecific anticholinergic drugs (e.g., scopolamine) that are used for treating vestibular conditions such as motion sickness. The effect of these drugs are only considered with respect to their central effects, such as suppression of neurons in the vestibular nuclei [91]. However, such peripheral changes as the source of inputs to the vestibular nuclei should also be taken into account.

An excitatory effect with inhibition of KCNQ channels similar to that observed in the present study has also been shown in peripheral Aδ fibers (in response to thermal or mechanical stimuli) [43] as well as pyramidal neurons in the CA1 region of the hippocampus [45, 92]. In the latter, KCNQ channels regulate the threshold [49] because of the colocalization of KCNQ channels with Na^+^ channels in the spike initiation zone of axons [47, 48]. As a result, inhibition of KCNQ channels in the CA1 region can produce spontaneous firing, even in the absence of synaptic inputs [49]. Similarly, KCNQ channels are being expressed near the AP initiation zone in calyces [31], which could explain the induction of spontaneous firing in some calyces during application of oxo-M or linopirdine.

### Conclusion

Our results suggest that cholinergic efferent inputs can modulate the activity of calyces and most likely irregular afferents through changes in the activity of KCNQ channels. Our findings suggest that without efferent inputs, open KCNQ channels result in low membrane resistances and high spike frequency adaptation, which precludes rapid firing of action potentials and most likely results in decreased excitability. Activation of mAChR by efferents and inhibition of KCNQ channels in calyces, result in lower thresholds, faster responses (i.e., decrease in first spike latency), and higher sensitivities (i.e., firing many APs during step depolarizations). In this way, efferents can play a role in modifying and adjusting the activity of calyx terminals and by extension, irregular afferents. Such adjustments could play a role in gauging responses to fast movements as required during daily activities or following changes in the peripheral sensory organs, such as loss of hair cells with aging.

## ACKNOWLEDGEMENTS

We thank Dr. Malcolm Slaughter and Dr. Elisabeth Glowatzki for comments on the manuscript and Dr. Matthew Johnson for helping in the early stages of setting up the experiments.

